# Introducing the digital PCR data essentials standard to harmonize data structure for clinical and research use

**DOI:** 10.64898/2026.04.10.717368

**Authors:** Trypsteen Wim, Vynck Matthijs, Untergrasser Andreas, Alexandra S Whale, Rödiger Stefan, Dobnik David, Bogožalec Košir Alexandra, Milavec Mojca, Kubista Mikael, Michael W Pfaffl, Abdel Nour Afif, Young-Kyung Bae, Stephen A Bustin, George A Calin, Chen Yao, Megan H Cleveland, Alfonso De Falco, Forootan Amin, Denise M O’Sullivan, Alison S Devonshire, Carole A Foy, Stephanie I Fraley, David G Gleerup, He Hua-Jun, Hellemans Jan, Lievens Antoon, Guro E Lind, Porco David, Romsos Erica, Thas Olivier, Drandi Daniela, Taly Valerie, Marie de Tayrac, Jim F Huggett, Vandesompele Jo, Ward De Spiegelaere

## Abstract

Digital PCR (dPCR) is a powerful technology for absolute quantification of nucleic acids, valued for its accuracy, sensitivity, and repeatability. Yet, the commercialization of different instruments with proprietary software has introduced challenges to data analysis, interoperability, and comparability. Therefore, we present the Digital PCR Data Essentials Standard (DDES) – a lightweight, human- and machine-readable, and cross-platform data standard developed in collaboration with the dPCR community. The standard consists of three file types designed to enable both manual inspection and automated analysis: (i) a *main* file summarizing experiment and reaction-level (meta-)data; (ii) an *assay* file describing targets and detection chemistry, and (iii) *intensity* files capturing partition-level raw fluorescence data per reaction. DDES supports a wide range of current dPCR applications, including singleplex and multiplex assays, endpoint and real-time readouts, and will be curated to implement future dPCR developments. By harmonizing the data structure, DDES lays out the foundation for FAIR dPCR data practices and supports improved software compatibility, collaborative and reproducible research, and future dPCR data repositories.

## Introduction

Digital PCR (dPCR) is a mature technology for rapid small-to medium-throughput absolute quantification of specific nucleic acid sequences (targets). It is based on subdividing the sample reaction mix into thousands of partitions and using the fluorescence read-out of these miniature PCR reactors as a binary readout to quantify the abundance of the intended target. Compared to conventional PCR techniques, dPCR is characterized by its high precision in quantifying low-abundance targets, enhanced sensitivity, and superior reproducibility [1-3]. Supported by early key research findings showing these advantages, together with a breadth of described applications, an increase in commercially available dPCR instruments has been observed over the last decade, confirming the popularity and broad usage of dPCR-based quantification [4-5].

Recent advances in dPCR have primarily focused on hardware improvements. Instrument providers are enhancing workflow integration and expanding core dPCR features, such as increasing the number of detectable fluorophores, boosting sample throughput, and offering random-access capabilities, as well as application-specific run formats with different numbers of partitions analyzed per sample. On the software side, manufacturers provide proprietary software with their instruments designed to support data acquisition and facilitate data analysis. However, as seen in fields such as “omics” technologies, these software solutions often do not support handling complex or customized experimental designs, sometimes exhibit subpar performance with limited end-user intervention, operate as a black-box, or lack specific downstream analysis options.

Besides the software development efforts of instrument providers, there is significant interest within the dPCR research community to develop third-party data analysis methods to resolve the above-mentioned shortcomings. These tools mainly focus on automation, thresholding, and data visualization (Table 1) [6-7]. As the field progresses, both third-party and proprietary software tools for the analysis of complex assays are expected to become available in the coming years. Such tools will be essential for expanding the utility of the dPCR technology, both in the context of increasingly sophisticated experimental designs of (higher-order) multiplexing, and its amenability to higher-throughput or regulatory settings [7-8].

**Table 1:** Overview of research-community developed dPCR data analysis software for partition classification. *\*NA: not applicable, implementation requires data preprocessing*.

| name | year | compatibility | reference |
| --- | --- | --- | --- |
| definetherain | 2014 | Bio-Rad QX100/QX200 | [11] |
| ddpcrquant | 2015 | Bio-Rad QX100/QX200<br>Stilla Naica<br>Thermo Fisher QuantStudio 3D | [12] |
| ddPCRmulti | 2015 | Bio-Rad QX100/QX200 | [13] |
| Cloudy | 2016 | NA* | [14] |
| ddpcr | 2016 | Bio-Rad QX series | [15] |
| Umbrella | 2017 | NA* | [16] |
| twoddpcr | 2017 | Bio-Rad QX series | [17] |
| Calico | 2017 | Bio-Rad QX100/QX200 | [18] |
| ddpcrclust | 2018 | Bio-Rad QX100/QX200 | [19] |
| PoDCall | 2023 | Bio-Rad QX series | [20] |
| Digital PCR<br>Cluster Predictor | 2023 | Bio-Rad QX series<br>Thermo Fisher Quantstudio 3D | [35] |
| dipcensR | 2024 | NA* | [8] |
| Polytect | 2025 | NA* | [21] |

Important drawbacks witnessed with both proprietary and third-party data analysis tools developed over the past decade, are a lack of (i) *interoperability, i*.*e*., both proprietary software and third-party data analysis tools are typically tailored to a single (or handful of) instrument(s) (Table 1), and (ii) *reproducibility*, as proprietary software is often closed-source and available only to those having purchased the instrument [6]. Moreover, new software releases or updates with modifications to data analysis pipelines may hamper repeatability of experiments and cause third-party software failures and compatibility issues due to subtle differences in the underlying data structures. These observations indicate an urgent need for a data standard compatible across dPCR instruments to mitigate these problems and ensure data consistency, as achieved for other technologies. For example, in the field of high-throughput sequencing (HTS), both commercial and third-party data analysis software are available. This ecosystem is enabled by a small set of open-source, cross-platform, widely accepted data standards (e.g., FASTQ and BAM), supporting adherence to FAIR data principles [9-10].

Similar data standards have been developed and are widely used elsewhere, such as for quantitative PCR (RDML [22] and RDES [23]), flow cytometry (FCS [24]) and mass spectrometry (mzML [25]) data. Most relevant to dPCR, the standard for quantitative PCR (qPCR) data was developed in 2009 and named ‘Real-time PCR Data Markup Language’ or RDML. RDML’s XML-based data files provide the raw qPCR data as well as relevant metadata to analyze the experiment, such as PCR cycling conditions, annotation and grouping of samples, and the combination of runs into one experiment [22]. Given the rather extensive scope of the RDML standard, the RDML consortium more recently developed a basic ‘Real-time PCR Data Essential Spreadsheet’ (RDES) format for exporting a single qPCR run, which contains only the essential annotation and the raw data of a qPCR experiment in a tabular format [23].

In contrast, no such data standard currently exists for dPCR, hampering the development of interoperable and reproducible data analysis tools, whether commercial or open source. Given the increasing data complexity, breadth of instruments and suppliers, and growing uptake of dPCR, the need for a standardized data format in dPCR is becoming increasingly urgent. For this purpose, the Digital PCR Data Essentials Standard (DDES) is introduced. DDES is a community-developed data standard supporting a wide range of current, emerging, and envisaged future dPCR experiment designs, including single-color singleplex, single- or multi-color multiplex, and higher-order multiplex experiments. The DDES structure is compatible with all current dPCR instruments, as well as emerging platforms, including instruments supporting real-time readout or high-resolution melting dPCR [26-27]. In line with the philosophy behind the RDES format, DDES is easily readable in any spreadsheet software or text file editor, owing to its table-like structure.

## Material and Methods

### Standard development

Authors WT, MV and WDS initiated a dPCR data standard working group at Ghent University in January 2023. After creating a crude outline for DDES, the preliminary data standard format was pitched to a network of experienced dPCR scientists at the 10th Gene Quantification Congress (March 20-24th 2023, Freising, Germany), and feedback gathered. In the second quarter of 2023, a revised format was distributed by e-mail to an extended network of experienced dPCR researchers, and further feedback gathered. Based on the comments received, the format was refined, and a pre-final version first presented to the public at the European Digital PCR Symposium 2024 (EUDIP2024, February 6th, 2024, Ghent, Belgium). Gathering and processing additional feedback, the data standard was further adapted and sent for a final review by expert dPCR users in July 2025. A final revision ultimately led to the version of DDES presented in this work.

### Data conversion

Data conversion software to parse dPCR data exported from commercial software was developed using the R/Shiny framework [28]. It is freely available through a web browser at https://digpcr.shinyapps.io/DDES/. This includes the extraction of DDES desired information from platform-specific data export formats.

## Results

### The Digital PCR Data Essentials Standard (DDES)

DDES is a universal dPCR data standard to store dPCR data generated by any instrument and is envisaged to be curated over time to adapt to new technological developments. It consists of three distinct file formats: the *main* file, *assay* file, and *intensity* file(s) and used DDES-related key terminology (Table 2 and 3). For each run, defined as a set of reactions wherein any combination of nucleic acid targets is measured, a DDES zip file is generated that, when unpacked, consists of (i) one *main* file, (ii) one *assay* file and (iii) several *intensity* files (one per reaction or well). Each file contains a header with metadata information, and the tabular dPCR data (Supplemental Data 1-3). Each of these files is designed to be both human- and machine-readable and is easily edited in a text editor or spreadsheet software. However, to avoid violation of the data format, it is mandatory to edit DDES files through a dedicated DDES-compatible interface. All three types of files adhere to a standardized file name structure, containing experiment and run identifiers, along with a date and time stamp. The latter allows for identifying iterations of the same experiment/run (Table 2).

**Table 2:**
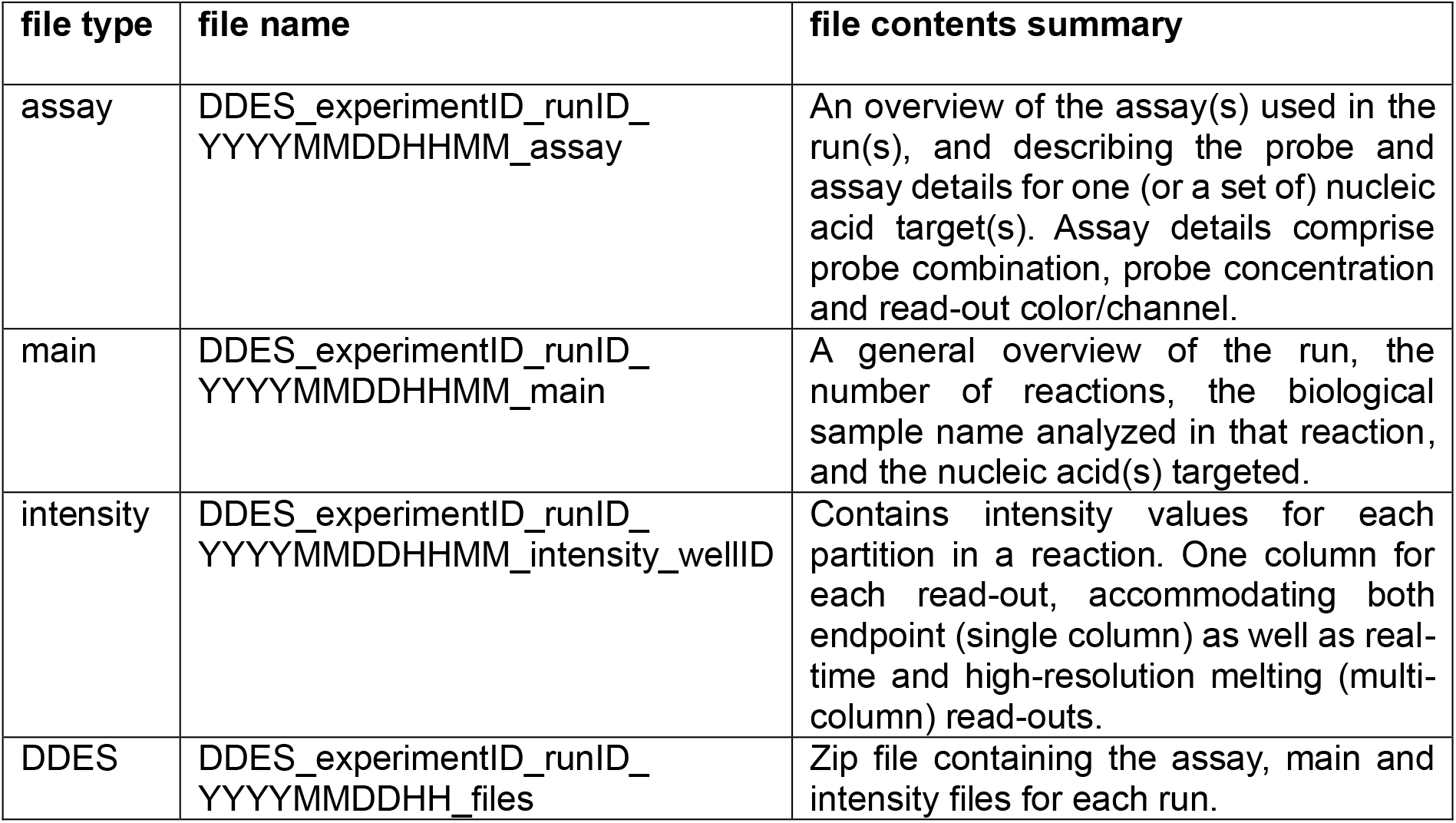
The three different file types of DDES. Their file names adhere to a given standard, and their file contents are well-defined.

| file type | file name | file contents summary |
| --- | --- | --- |
| assay | DDES_experimentID_runID_YYYYMMDDHHMM_assay | An overview of the assay(s) used in the run(s), and describing the probe and assay details for one (or a set of) nucleic acid target(s). Assay details comprise probe combination, probe concentration and read-out color/channel. |
| main | DDES_experimentID_runID_YYYYMMDDHHMM_main | A general overview of the run, the number of reactions, the biological sample name analyzed in that reaction, and the nucleic acid(s) targeted. |
| intensity | DDES_experimentID_runID_YYYYMMDDHHMM_intensity_wellID | Contains intensity values for each partition in a reaction. One column for each read-out, accommodating both endpoint (single column) as well as real-time and high-resolution melting (multi-column) read-outs. |
| DDES | DDES_experimentID_runID_YYYYMMDDHH_files | Zip file containing the assay, main and intensity files for each run. |

**Table 3:** DDES related terms and definitions.

| Key Term | Definition |
| --- | --- |
| Target | A specific nucleic acid sequence of interest that is amplified and quantified with digital PCR. |
| Run | A group of wells (plate or strip) that undergo a certain dPCR reaction. Depending on the platform these are grouped per dPCR consumable (plate/chip) that undergo the same cycling protocol. |
| Replicate measurements | A replicate reaction is considered when performing the same dPCR reaction on input material originating from the same sample stock but in different wells or tubes. |
| Experiment | An experiment can consist of one or multiple dPCR runs. DDES will export information at the run-level, meaning the experimentID can be shared among DDES exports but will have different runIDs linked to it. |
| Well, Sample or dPCR reaction | All components that make up the analysis of a single well or tube related to the dPCR reaction: sample-specific PCR mix, partitioning, cycling protocol and imaging. |
| Assay | One or multiple PCR reaction(s) that quantify a certain target inside the same dPCR reaction. |

The first of three file types is the “*main*” file (Supplemental Data 1). It is the ‘informational heart’ of a dPCR run and contains a tabular overview of all reactions. For each reaction, the *main* file contains information about the target(s) and sample(s) analyzed, while linking to the *assay* and *intensity* files via the assay ID and well ID, respectively. Based on the assay ID that is linked to the sample, each target that is measured is represented on a separate line – so that the data for a triplex assay would cover three lines giving the target ID (different for the three lines), assay ID (same for the three lines) with the number of positive and negative partitions and concentration estimate for the target. For targets detected with a single color, a threshold value can be included. The *main* file can stand alone for subsequent basic data analysis such as absolute or relative concentration estimation.

The second file type is the “*assay*” file (Supplemental Data 2). An *assay* file contains information on all assays used in a run; each assay has a unique assay ID and is defined as any combination of nucleic acid targets linked to their respective color or channel detection strategy. Such information for each assay includes (i) the target name(s), (ii) the read-out channel(s) for each target, and (iii) the probe reaction concentrations, if applicable. Each run is associated with a single *assay* file generated in the DDES data package (Figure 1).

**Figure 1.**
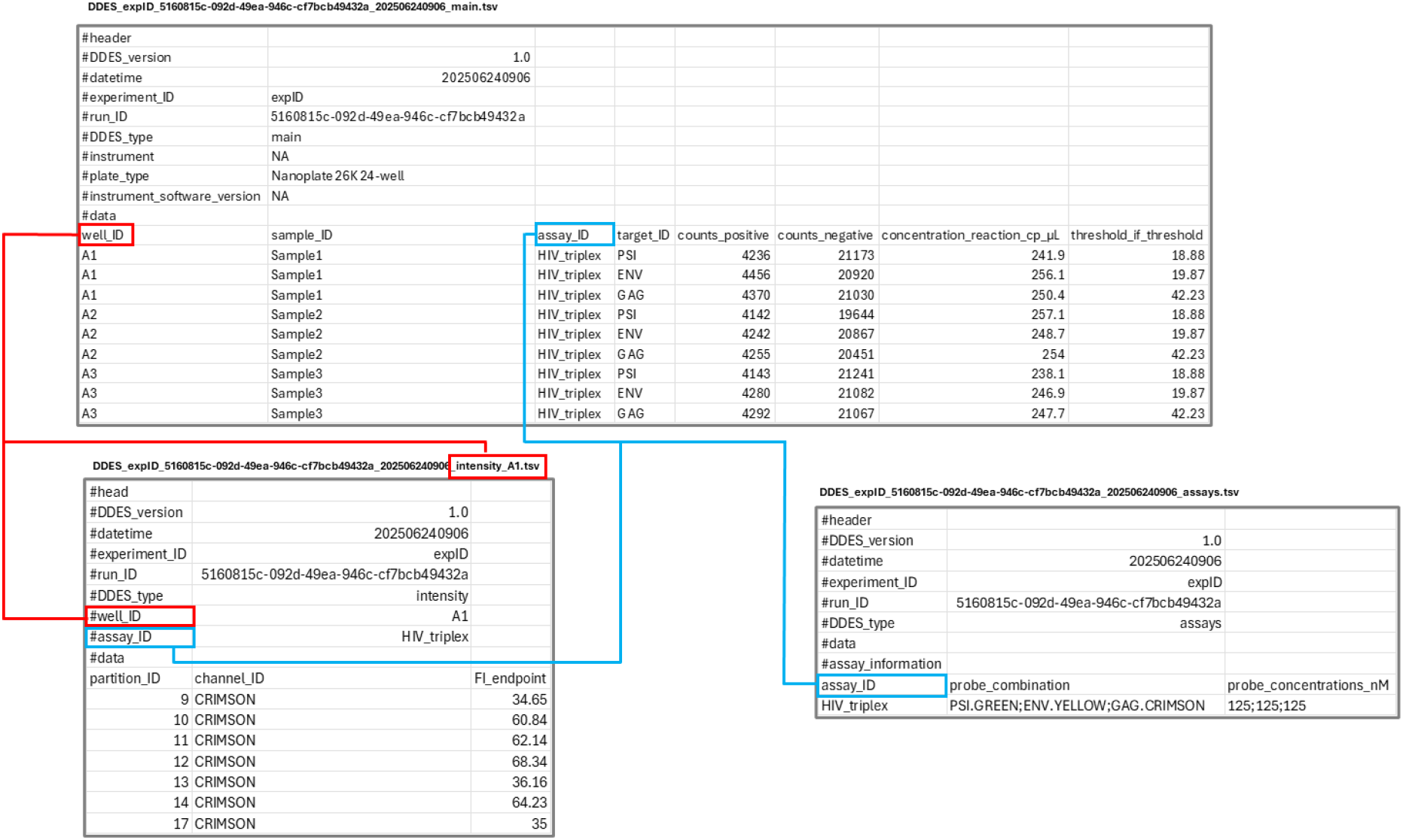
Illustration of the DDES data package for each run and linking of the files through the well ID and assay ID. The *main* file consists of the summary information of the run and is linked via the well ID to the *intensity* files to retrieve fluorescence values of the partitions for that well. The well ID is also used to link the *main* file with the *assay* file to retrieve the corresponding assay ID for that well ID. This example is a data set exported from the QIAcuity Four instrument and contains information from three reactions containing a triplex assay which measured 3 targets with one color per target.

The third, and final file type, is the “*intensity*” file (Supplemental Data 3). Each run will typically entail multiple *intensity* files: one per reaction (*i*.*e*., as opposed to a single *main* and *assay* file). The *intensity* files contain a single- or multi-column data section, listing the endpoint or real-time cycle fluorescence read-out (column(s)) for all valid partitions (rows) in a reaction. The intensity values included in the DDES format are the instrument-reported partition intensities after proprietary processing steps by the instrument. In addition, DDES intensity files only contain valid partitions that pass the instrument and/or end-user quality control on the instrument software itself; the instrument software is expected to prefilter platform exports that include quality control measurement information for each partition. For multi-color experiments, a partition will return multiple rows, one for each detection channel. To accommodate data from platforms that have real-time or high-resolution melting capabilities, the fluorescence values can be presented in multiple columns for a given partition and color, in the same philosophy as RDES does for qPCR data.

Finally, each file contains a minimal metadata header showing information related to the DDES version and the run, experiment or reaction itself, such as the type of instrument, run ID, consumable, and software version used for generating the read-out (see Supplemental Data 1-4 for a complete and detailed description).

### Converting disparate dPCR data formats to DDES

While we encourage dPCR instrument vendors to export directly in DDES format, we have developed a data conversion software framework to parse dPCR data exported from instrument software using the R/Shiny framework (Supplemental Data 4) [28]. For information that is not available in the platform exports, the DDES data structure will report ‘NA’ (Not Available) values. As an example, using the QIAcuity export data formats from a 3-color HIV-1 DNA data set, a DDES converter is freely available through a web browser interface at https://digpcr.shinyapps.io/DDES/ and currently supports QIAcuity and QX200 data format conversion. In addition, source code is made available at the DIGPCR Center GitHub page, supportive of a broad adoption and curation of the converter by the dPCR community. This ongoing collaborative approach between the DDES consortium and dPCR community to curate the DDES converter for compatibility with dPCR platforms increases the likelihood of remaining current with new platforms and software developments.

## Discussion

DDES is minimal, containing only essential dPCR data, and designed to be compatible with all currently available dPCR instruments, with room to accommodate future instrument developments. For example, higher-order multiplexed assays, such as color combination, where two (or more) colors per target are used, or amplitude multiplexing, where two (or more) targets per color are used, are still in their infancy. Their advantages are numerous, and early studies suggest they will become increasingly common. Similarly, moving beyond endpoint readouts to incorporate real-time detection, as routinely done in qPCR, is expected to become more widespread in dPCR. To ensure that DDES is a robust, future-proof standard, compatible with the broad field of current and emerging dPCR applications, it has been designed in collaboration with, and backed by, the dPCR user community. The maintenance and further development of DDES will be governed by the DIGPCR Center, in consultation with the scientific community. In addition, DDES is also made applicable in the context of clinical laboratories reporting on patient test results. DDES does not require personal identifiers but implemented user-defined sample IDs that can be linked to any clinical metadata outside of the DDES framework, adhering to privacy regulations.

DDES is not intended to share all unprocessed raw measurements. Depending on the dPCR instrument, DDES data will undergo some pre-processing, such as quality control or color-compensation. The focus of DDES is to provide only the essential data needed for exchanging dPCR results, independently from the instrument that was used to generate them. We believe that this will improve the reproducibility of dPCR experiments as, by sharing data in a universal format, DDES makes it easier to assess data quality through compatibility with any laboratory data analysis pipeline.

In terms of absolute quantification based on DDES formatted information, we urge the platform providers to be transparent regarding their calculations and disclose (average) partition volumes. Using the DDES output, lambda values and total copies per reaction with their uncertainty can be (re)calculated in a reproducible manner. However, variability may be introduced if different assumptions are made regarding single partition volume or the total partition volume (*i*.*e*., the cycled volume) of a reaction. As for the uncertainty or variance estimation and alongside calculations on the quantification result, we emphasize using dedicated generic statistical frameworks or tools to adequately estimate the total variation introduced by the dPCR workflow and experimental setup (e.g. replicate measurements, copy number variation or fraction abundance), as reporting Poisson error estimates are overoptimistic in reporting precision or confidence intervals (29).

Improving reproducibility is a key objective of the “Minimum Information for Publication of Quantitative Digital PCR Experiments” (dMIQE). While DDES and dMIQE share this common goal, DDES and dMIQE are complementary: DDES does not intend to replace dMIQE, but to strengthen its objectives. To this end, overlapping information between dMIQE and DDES is kept to a bare minimum, with duplicated info in DDES only retained when necessary for proper data analysis. As well as improving the quality of the published methods, the DDES standard file format could also support the application of dPCR for nucleic acid reference material value assignment [30]. There is a growing interest in digitization of certified reference material certificates, with linked availability of the data corresponding to the reference material characterization studies.

The minimalistic approach chosen for DDES is deliberate. The addition of several features was contemplated but deemed to decrease its ease of adoption, inter-instrument and inter-assay compatibility, and human readability. Therefore, DDES mainly focuses on limited but mandatory (meta-)data fields, uniform data storage, readability, and minimal data duplication. Storing information in multiple files slightly complicates data exchange, but is tackled with a single zipped data package folder for each experiment. This approach contrasts with other data standards that often store data in a single file per experiment or reaction (*e*.*g*., sequencing’s FASTQ or mass spectrometry’s mzML file format), arguably, at the expense of human-readability. Human-readable files are often preferred, as exemplified by the development of RDES after the more elaborate, but more complicated RDML format [23].

Last, while DDES is a lightweight data framework that is easily shared, dPCR’s data FAIRness (Findable, Accessible, Interoperable, Reusable) would benefit from a dedicated, searchable dPCR repository, in line with such repositories that are available for sequencing (*e*.*g*., Sequence Read Archive, European Nucleotide Archive [31-32]), real-time quantitative PCR (*e*.*g*., RDMLdb [33]), or mass spectrometry (*e*.*g*., MassBank [34]) data. The combined storage of dMIQE information and DDES data in such a repository could further increase dPCR’s data FAIRness. Further, the use of a controlled vocabulary for describing, *e*.*g*., sample and assay types would enhance dPCR’s data FAIRness; however, due to the number of possibilities and nuances, an agreement was, for now, not reached among the author group. This remains an important area of development for future DDES versions.

In conclusion, DDES addresses a critical and growing gap in the dPCR ecosystem by providing a universal, interoperable, and minimal data format tailored to the needs of both researchers and software developers. By prioritizing simplicity, transparency, and cross-platform compatibility, DDES lays out the groundwork for more reproducible, scalable and collaborative dPCR research. As adoption grows, DDES will not only facilitate data exchange and analysis, but also support the development of robust, platform-independent software tools. Looking ahead, its integration into community standards, journals, and data repositories will be key to establishing FAIR principles in dPCR workflows. Ultimately, DDES contributes to making dPCR a widely accessible and reproducible technology for molecular quantification.

## Abbreviations

dPCR: digital polymerase chain reaction
HTS: High-Throughput Sequencing
RDML: Real-time PCR Data Markup Language
RDES: Real-time PCR Data Essential Spreadsheet
FCS: flow cytometry standard
qPCR: quantitative PCR
DDES: Digital PCR Data Essentials Standard
dMIQE: Minimum Information for Publication of Quantitative Digital PCR Experiments.

## Data Availability

The dPCR data used in the Shiny web application is available through Github (www.github.com/digpcr/DDES).

## Code Availability

The code for the Shiny web application of the DDES converter is available through GitHub (www.github.com/digpcr/DDES). The DDES shiny web application is available through the DIGPCR shinyapps server at https://digpcr.shinyapps.io/DDES.

## Author Contributions

WT,WDS,MV,OT,JFH,ASW,AU,SR contributed to conceptualization. WT,MV have developed the data conversion tool. WT,WDS,MV have written the manuscript. All authors contributed to reviewing, editing and proofing of the manuscript. WT,WDS are responsible for coordinating and communicating with all the authors.

## Acknowledgments

WT, MV, JV and WD gratefully acknowledge the DIGPCR Core facility at Ghent University (Belgium) for the use and support on digital PCR. MK acknowledges the project MULTIOMICS_CZ (Program Johannes Amos Comenius, Ministry of Education, Youth and Sports of the Czech Republic, ID Project CZ.02.01.01/00/23_020/0008540) – Co-funded by the European Union and 86652036 from RVO. Points of view in this document are those of the authors and do not necessarily represent the official position or policies of the U.S. Department of Commerce or the Department of Justice. Certain commercial equipment, instruments, and materials are identified in order to specify experimental procedures as completely as possible. In no case does such identification imply a recommendation or endorsement by NIST, nor does it imply that any of the materials, instruments, or equipment identified are necessarily the best available for the purpose.

