## Supplemental Data for "Introducing the digital PCR data essentials standard to harmonize data structure for clinical and research use"

### Supplemental Data (Figures, Tables, Methods)

### Supplemental Data 1: DDES *main* file

### Supplemental Data 1A: DDES *main* file (Example)

**DDES_expID_5160815c-092d-49ea-946c-cf7bcb49432a_202506240906_main.tsv**

| | | | | | | | |
|---|---|---|---|---|---|---|---|
| #header | | | | | | | |
| #DDES_version | 1.0 | | | | | | |
| #datetime | 202506240906 | | | | | | |
| #experiment_ID | expID | | | | | | |
| #run_ID | 5160815c-092d-49ea-946c-cf7bcb49432a | | | | | | |
| #DDES_type | main | | | | | | |
| #instrument | NA | | | | | | |
| #plate_type | Nanoplate 26K 24-well | | | | | | |
| #instrument_software_version | NA | | | | | | |
| #data | | | | | | | |
| well_ID | sample_ID | assay_ID | target_ID | counts_positive | counts_negative | concentration_reaction_cp_µL | threshold_if_threshold |
| A1 | Sample1 | HIV_triplex | PSI | 4236 | 21173 | 241.9 | 18.88 |
| A1 | Sample1 | HIV_triplex | ENV | 4456 | 20920 | 256.1 | 19.87 |
| A1 | Sample1 | HIV_triplex | GAG | 4370 | 21030 | 250.4 | 42.23 |
| A2 | Sample2 | HIV_triplex | PSI | 4142 | 19644 | 257.1 | 18.88 |
| A2 | Sample2 | HIV_triplex | ENV | 4242 | 20867 | 248.7 | 19.87 |
| A2 | Sample2 | HIV_triplex | GAG | 4255 | 20451 | 254 | 42.23 |
| A3 | Sample3 | HIV_triplex | PSI | 4143 | 21241 | 238.1 | 18.88 |
| A3 | Sample3 | HIV_triplex | ENV | 4280 | 21082 | 246.9 | 19.87 |
| A3 | Sample3 | HIV_triplex | GAG | 4292 | 21067 | 247.7 | 42.23 |

**Supplemental Data 1B: DDES *main* file (Data Schema)**


```
{
  "file_format": {
    "extension": "tsv"
    "separator": tab,
    "decimal": ".",
    "missing_value": "NA",
    "metadata_prefix": "#",
    "metadata_marker": "#head",
    "main_marker": "#main"
  },
  "metadata_fields": [
    {
      "name": "DDES_version",
      "type": "character",
      "description": "Version of DDES standard",
      "restriction": none,
      "example": "1.0.0"
    },
    {
      "name": "datetime",
      "type": "character",
      "description": "Date and time of creation/change, YYYYMMDDHHMM format",
      "restriction": none,
      "example": "201212312359"
    },
    {
      "name": "experiment_ID",
      "type": "character",
      "description": "A user-defined experiment ID",
      "restriction": none,
      "example": "TriplexEGFR"
    },
    {
      "name": "run_ID",
      "type": "character",
      "description": "A user-defined run ID",
      "restriction": none,
      "example": "OptimPrimerConc"
    },
    {
      "name": "DDES_type",
      "type": "character",
      "description": "One of three DDES types: main, intensity, asssay",
      "restriction": none,
      "example": "main"
    },
    {
      "name": "instrument",
      "type": "character",
      "description": "Instrument description",
      "restriction": none,
      "example": "NioE1234"
    },
    {
      "name": "plate_type",
      "type": "character",
      "description": "Plate description",
      "restriction": none,
      "example": "Ruby20250131"
    },
    {
      "name": "instrument_software_version",
      "type": "character",
      "description": "Version of the instrument software used",
      "restriction": none,
```

```
      "example": "1.1.0"
    }
  ],
  "columns": [
    {
      "name": "well_ID",
      "type": "character",
      "description": "Well identifier",
      "restriction": none,
      "required": true,
      "example": "A01"
    },
    {
      "name": "sample_ID",
      "type": "character",
      "description": "Sample identifier",
      "restriction": none,
      "required": true,
      "example": "PosCtrl1"
    },
    {
      "name": "assay_ID",
      "type": "character",
      "description": "Assay identifier",
      "restriction": "Should occur in the assay file column assay_ID",
      "required": true,
      "example": "Assay1"
    },
    {
      "name": "target_ID",
      "type": "integer",
      "description": "Target identifier",
      "restriction": "Should occur in the assay file column probe_combination",
      "required": true,
      "example": 1
    },
    {
      "name": "counts_positive",
      "type": "integer",
      "description": "Number of positive partitions",
      "restriction": none,
      "required": true,
      "example": 15684
    },
    {
      "name": "counts_negative",
      "type": "integer",
      "description": "Number of negative partitions",
      "restriction": none,
      "required": true,
      "example": 2397
    },
    {
      "name": "threshold_if_threshold",
      "type": "double",
      "description": "Threshold, if a linear threshold was used",
      "restriction": none,
      "required": true,
      "example": 5842.4
    }
  ]
}
```

**Supplemental Data 2: DDES *assay* file**

**Supplemental Data 2A: DDES *assay* file (example)**

**DDES_expID_5160815c-092d-49ea-946c-cf7bcb49432a_202506240906_assays.tsv**

| | | |
|---|---|---|
| #header | | |
| #DDES_version | 1.0 | |
| #datetime | 202506240906 | |
| #experiment_ID | expID | |
| #run_ID | 5160815c-092d-49ea-946c-cf7bcb49432a | |
| #DDES_type | assays | |
| #data | | |
| #assay_information | | |
| assay_ID | probe_combination | probe_concentrations_nM |
| HIV_triplex | PSI.GREEN;ENV.YELLOW;GAG.CRIMSON | 125;125;125 |

### Supplemental Data 2B: DDES *assay* file (Data Schema)


```
{
 "file_format": {
  "extension": "tsv"
  "separator": tab,
  "decimal": ".",
  "missing_value": "NA",
  "metadata_prefix": "#",
  "metadata_marker": "#header",
  "main_marker": "#assay_information"
 },
 "metadata_fields": [
  {
   "name": "DDES_version",
   "type": "character",
   "description": "Version of DDES standard",
   "restriction": none,
   "example": "1.0.0"
  },
  {
   "name": "datetime",
   "type": "character",
   "description": "Date and time of creation/change, YYYYMMDDHHMM format",
   "restriction": none,
   "example": "201212312359"
  },
  {
   "name": "experiment_ID",
   "type": "character",
   "description": "A user-defined experiment ID",
   "restriction": none,
   "example": "TriplexEGFR"
  },
  {
   "name": "run_ID",
   "type": "character",
   "description": "A user-defined run ID",
   "restriction": none,
   "example": "OptimPrimerConc"
  },
  {
   "name": "DDES_type",
   "type": "character",
   "description": "One of three DDES types: main, intensity, asssay",
   "restriction": none,
   "example": "assay"
  }
 ],
 "columns": [
  {
   "name": "assay_ID",
   "type": "character",
   "description": "Assay identifier",
   "restriction": none,
   "required": true,
   "example": "Assay1"
  },
  {
   "name": "probe_combination",
   "type": "character",
   "description": "Combination of probes used in the assay, in the format 'target_ID.channel_ID'; if multiple, separated by a semicolon",
   "restriction": none,
   "required": true,
   "example": "1.1;1.2;2.1"
  },
```

```
    {
      "name": "probe_concentrations_nM",
      "type": "character",
      "description": "Concentration of probe(s) used, if multiple, separated by a semicolon",
      "restriction": none,
      "required": true,
      "example": "125;125;250"
    }
  ]
}
```

**Supplemental Data 3: DDES *intensity* file**

**Supplemental Data 3A: DDES *intensity* file (example)**

**DDES_expID_5160815c-092d-49ea-946c-cf7bcb49432a_202506240906_intensity_A1.tsv**

| | | |
|---|---|---|
| #head | | |
| #DDES_version | 1.0 | |
| #datetime | 202506240906 | |
| #experiment_ID | expID | |
| #run_ID | 5160815c-092d-49ea-946c-cf7bcb49432a | |
| #DDES_type | intensity | |
| #well_ID | A1 | |
| #assay_ID | HIV_triplex | |
| #data | | |
| partition_ID | channel_ID | FI_endpoint |
| 9 | CRIMSON | 34.65 |
| 10 | CRIMSON | 60.84 |
| 11 | CRIMSON | 62.14 |
| 12 | CRIMSON | 68.34 |
| 13 | CRIMSON | 36.16 |
| 14 | CRIMSON | 64.23 |
| 17 | CRIMSON | 35 |

**Supplemental Data 3B: DDES *intensity* file (Data Schema)**


```
{
 "file_format": {
   "extension": "tsv"
   "separator": tab,
   "decimal": ".",
   "missing_value": "NA",
   "metadata_prefix": "#",
   "metadata_marker": "#head",
   "main_marker": "#data"
 },
 "metadata_fields": [
   {
     "name": "DDES_version",
     "type": "character",
     "description": "Version of DDES standard",
     "restriction": none,
     "example": "1.0.0"
   },
   {
     "name": "datetime",
     "type": "character",
     "description": "Date and time of creation/change, YYYYMMDDHHMM format",
     "restriction": none,
     "example": "201212312359"
   },
   {
     "name": "experiment_ID",
     "type": "character",
     "description": "A user-defined experiment ID",
     "restriction": none,
     "example": "TriplexEGFR"
   },
   {
     "name": "run_ID",
     "type": "character",
     "description": "A user-defined run ID",
     "restriction": none,
     "example": "OptimPrimerConc"
   },
   {
     "name": "DDES_type",
     "type": "character",
     "description": "One of three DDES types: main, intensity, asssay",
     "restriction": none,
     "example": "intensity"
   },
   {
     "name": "well_ID",
     "type": "character",
     "description": "Well identifier",
     "restriction": none,
     "example": "A01"
   },
   {
     "name": "assay_ID",
     "type": "character",
     "description": "Assay identifier",
     "restriction": "Should occur in the assay file column assay_ID",
     "example": "Assay1"
   }
 ],
 "columns": [
   {
     "name": "partition_ID",
     "type": "integer",
```

```
      "description": "Partition identifier",
      "restriction": none,
      "required": true,
      "example": 1
    },
    {
      "name": "channel_ID",
      "type": "integer",
      "description": "Channel identifier",
      "restriction": none,
      "required": true,
      "example": 1
    },
    {
      "name": "FI_endpoint",
      "type": "double",
      "description": "Endpoint fluorescence intensity",
      "restriction": none,
      "required": true,
      "example": 8421.6
    },
    {
      "name": "FI_cycle1",
      "type": "double",
      "description": "Fluorescence intensity after 1 PCR cycle",
      "restriction": none,
      "required": false,
      "example": 100.1
    },
    {
      "name": "FI_cycleX",
      "type": "double",
      "description": "Fluorescence intensity after X PCR cycles (as many columns can be used as cycles
conducted)",
      "restriction": none,
      "required": false,
      "example": 200.1
    }
  ]
}
```

**Supplemental Data 4: The DDES web application**

**4A. DDES web tool - start page.** On the left you have the option of either selecting a "Demo Data set" to browse the DDES format or Upload a Data Set for DDES Conversion.

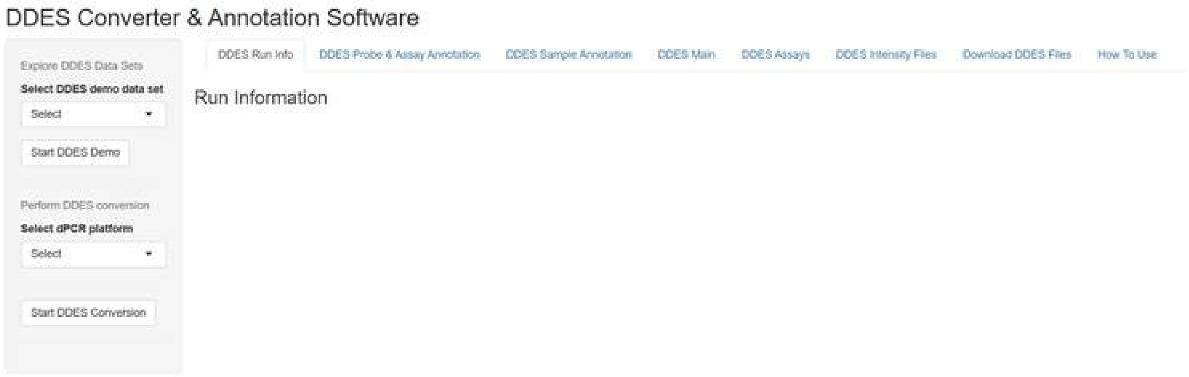


**4B. DDES web tool - selecting and uploading data set.** On the left you select the dPCR platform, which will trigger platform-specific file upload requirements. These are files you need to extract from the platform of the instrument provider. For the QIAcuity example, this requires the Analysis, RFU and plate layout file. Once clicked on "Start DDES Conversion", the software will retrieve and restructure your data set into the DDES format.

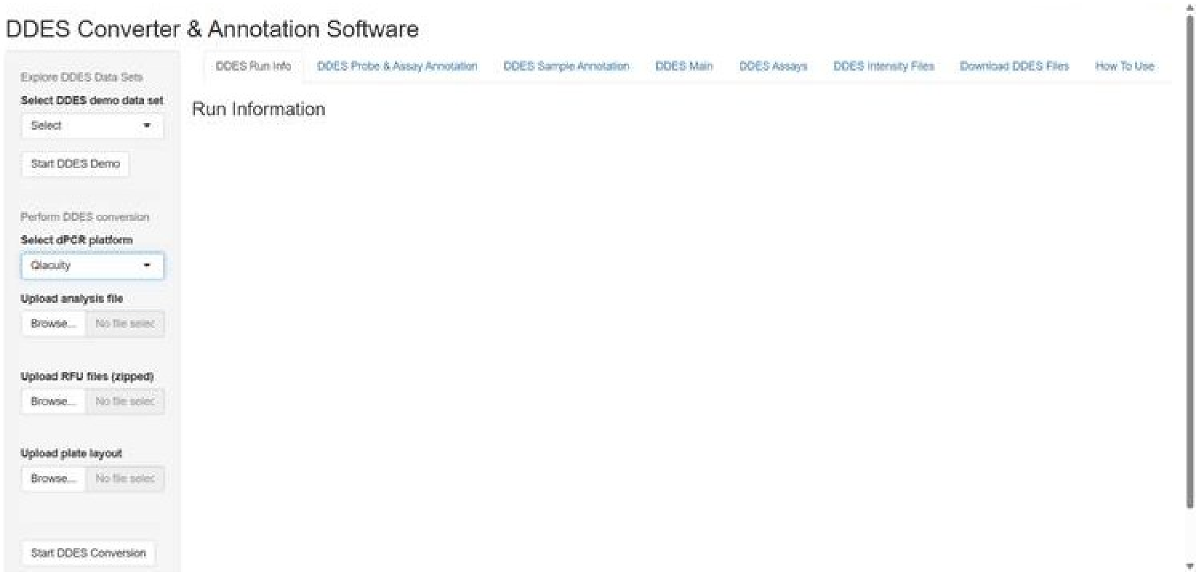

**4C. DDES web tool - run info tab.** Here minimal metadata is retrieved from the uploaded files. If this info is not available, the end user can provide this information or update the existing info.

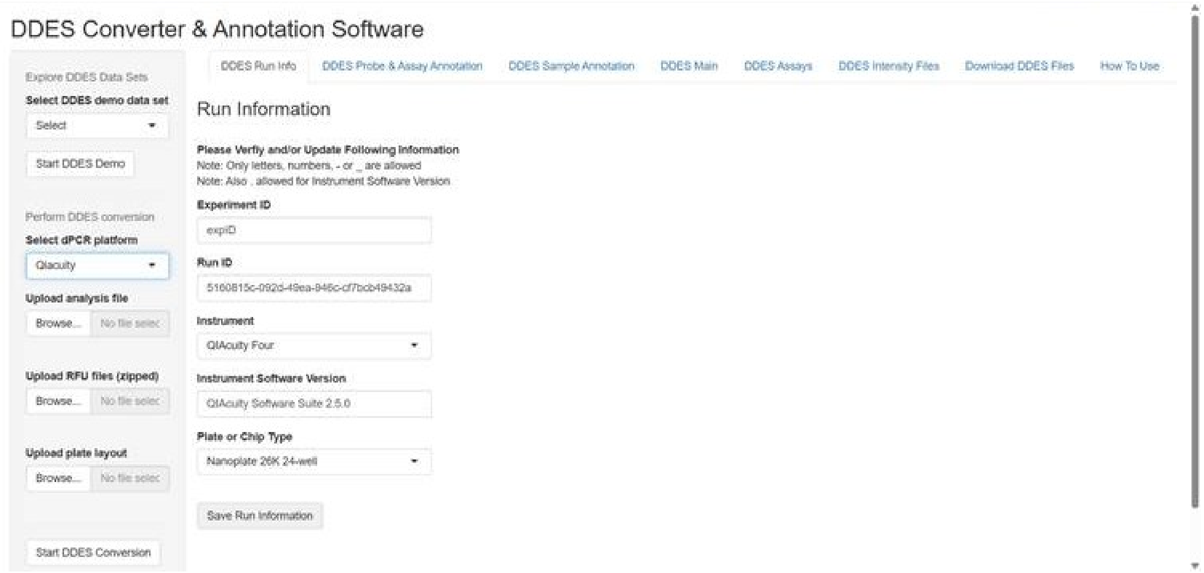


**4D. DDES web tool - probe and assay annotation tab.** Probes and Assay information is retrieved when available. The user can change or add probes in the probe table by making use of the add/delete buttons and providing probe names and link the appropriate channels (via a dropdown). The selector column can be used to delete probes. After annotation the user needs to click the save probe info button to lock the changes.

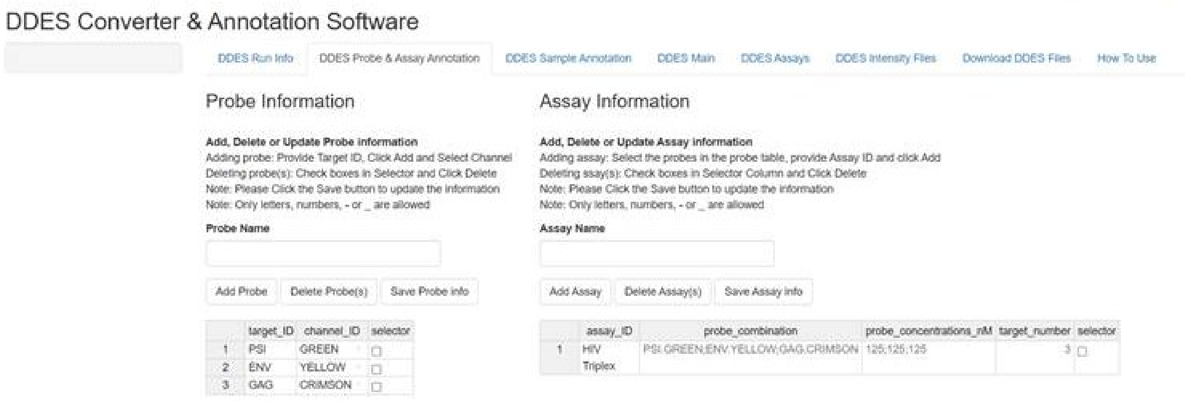

**4E. DDES web tool – probe and assay annotation tab.** The user can change or add assays in the assay table. Adding assays can be done by typing in assay name and click the Add Assay button. Next, the user can click the probe_combination column to trigger the probe selection for that assay. Next, the user can optionally click the probe_concentrations_nM column to provide the used probe concentration values. Finally, the user needs to click the Save Assay Info button to store the changes.

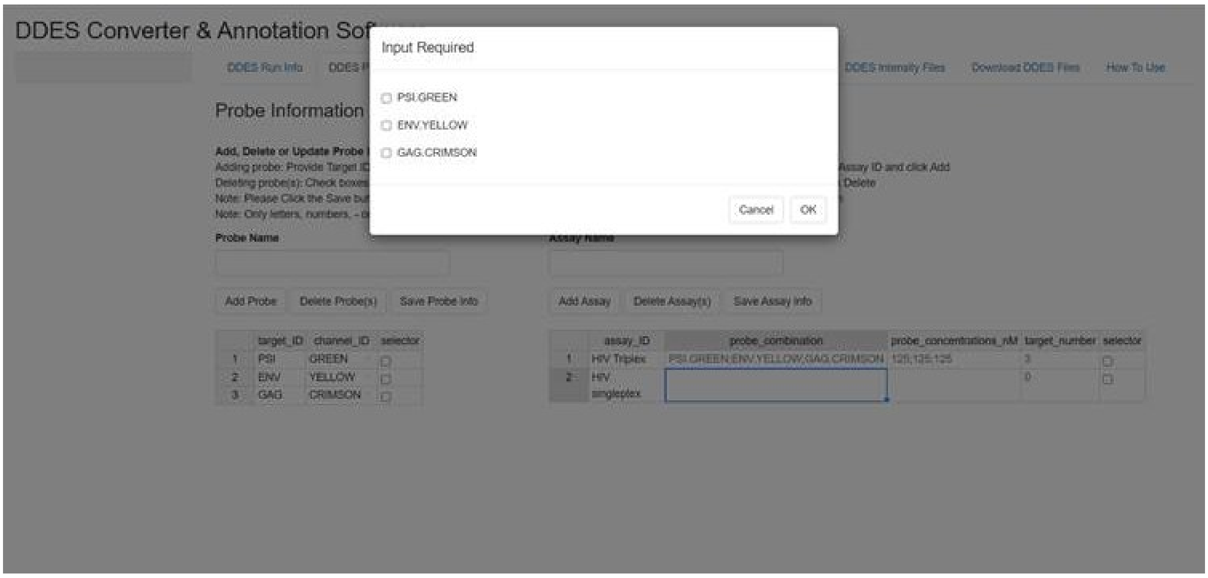


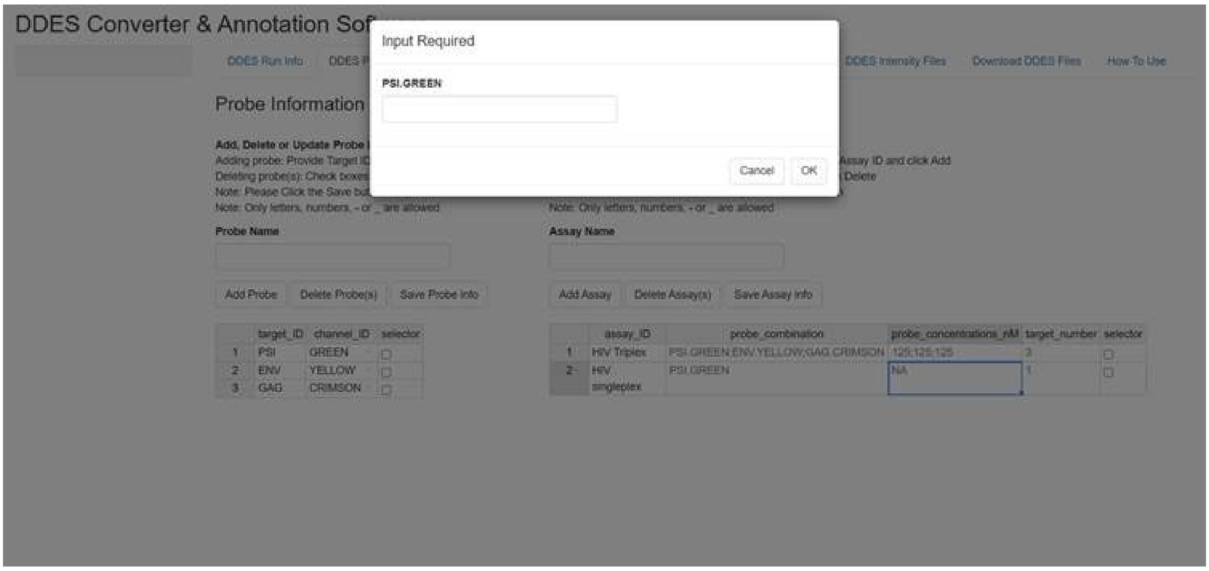

**4F. DDES web tool - sample annotation tab.** This tab shows information on the wells/samples found in the uploaded data set. A well_ID, assay_ID and sample_ID is retrieved and is editable by the user. By using the selector column the user can change the assay used via the dropdown assay selection menu. In the sample_ID column the user can copy paste or type in the sample IDs.

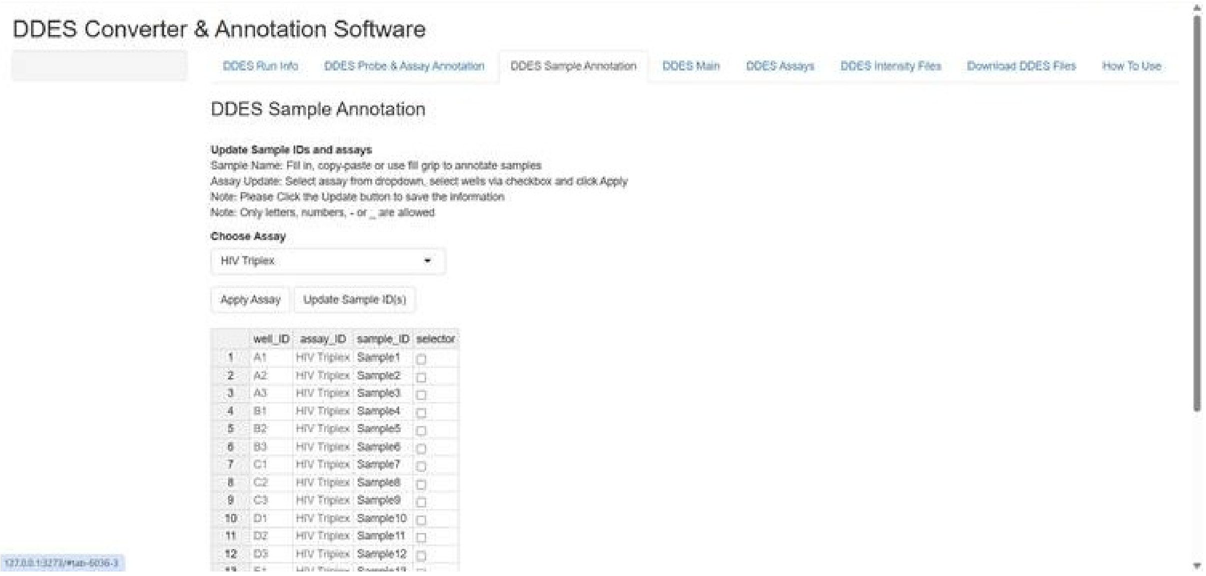


**4G. DDES web tool - main overview tab.** Tab for viewing the DDES main file.

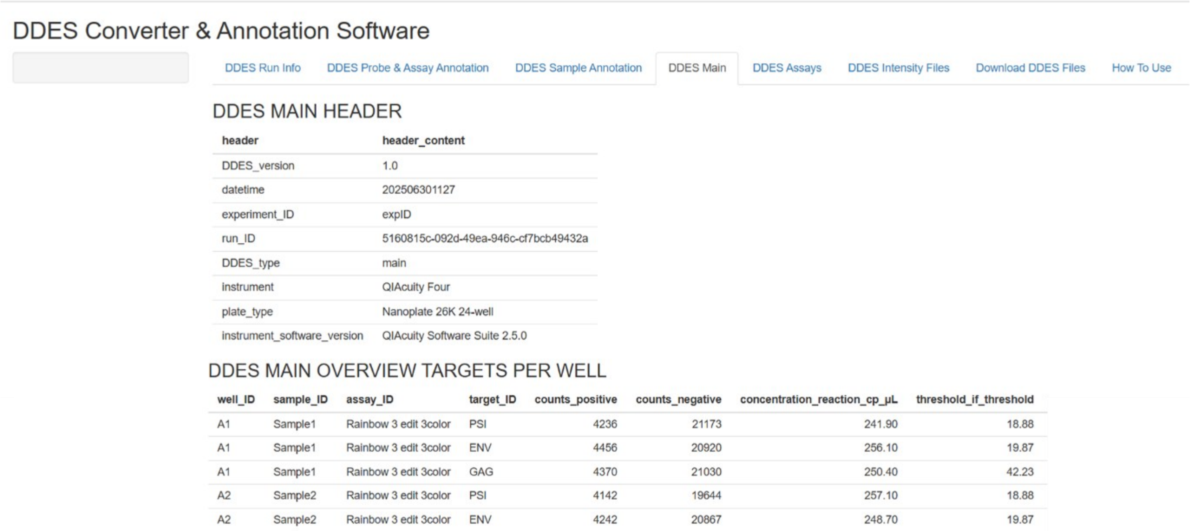

**4H. DDES web tool - assay overview tab.** Tab for viewing the DDES assays file.

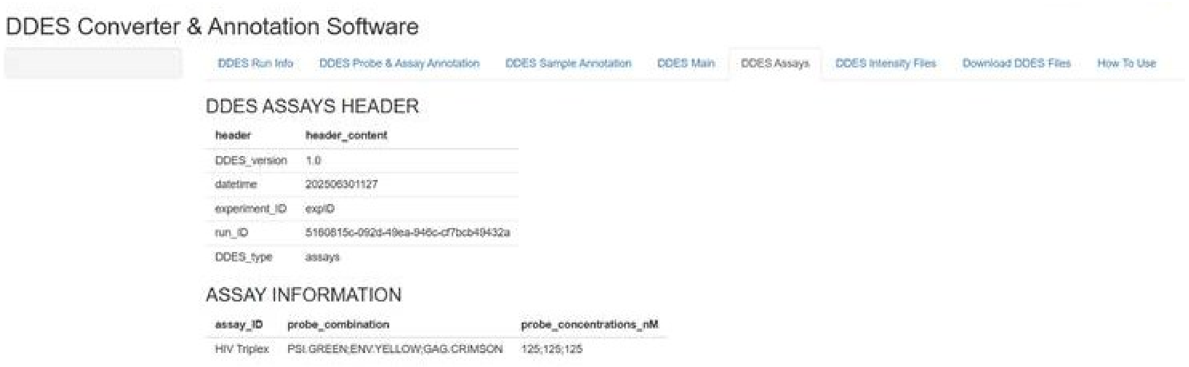


**4I. DDES web tool - intensity file tab.** Tab for viewing the DDES Intensity files.

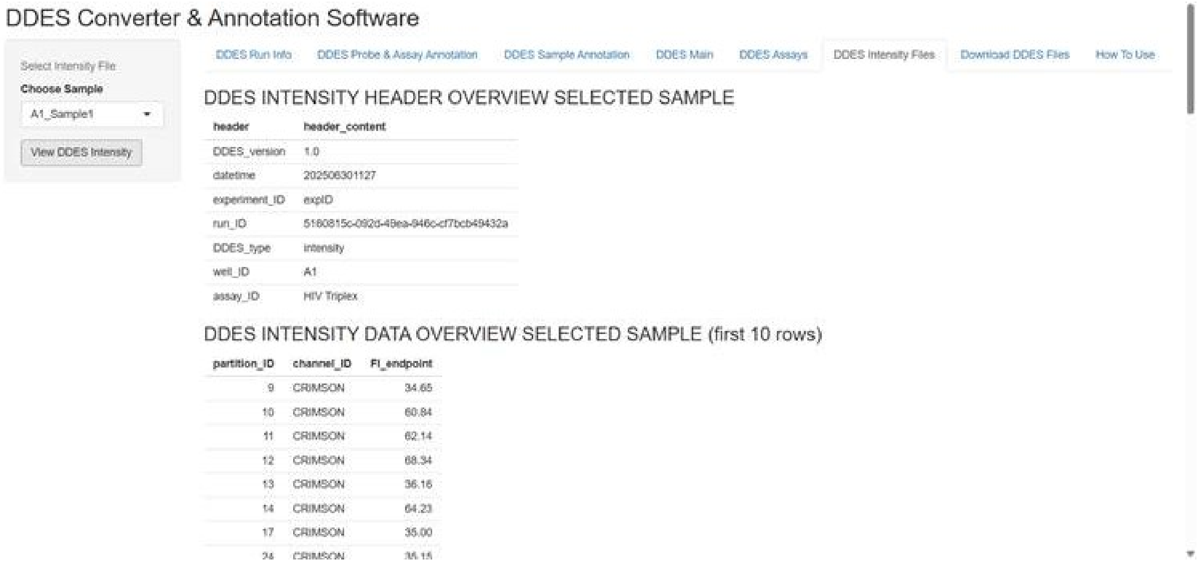

**4J. DDES web tool - download DDES file tab.** Here, the user can download the DDES files as a single zip folder.

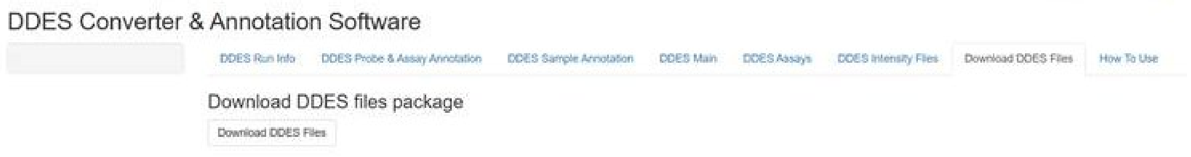
